# Analyzing an organism’s sensors using Maximum Entropy models with bias, variance, and confusion matrices

**DOI:** 10.64898/2026.01.12.698979

**Authors:** Christopher Wang, Elianna Schimke, Tristan Kako, Aiden Gao, Martina Lamberti, Joost le Feber, Sarah Marzen

## Abstract

Biological organisms have sensors that communicate information about the environment. Analyzing how well these biological sensors function has usually been done with mutual information between the sensor signal and the environment, but that can be computationally intractable and summarize something quite complex with just a single number. We suggest that alternatively, one may profitably analyze these biosensors using bias and variance or confusion matrices, depending on the kind of environment. Stimulus-dependent Maximum Entropy models are used to develop estimators of the environmental state given the sensor state, and these estimators in turn are then used to calculate either the bias and variance of the estimator or confusion matrices. We focus on several examples to understand the utility of non-information-based analyses: ligand-receptor binding models spanning genetic regulation to neuronal communication to bacterial chemotaxis, and spin-glass Ising models for neural activity in cultured neurons. These new computationally-efficient analyses add insight to existing analyses based on mutual information; in particular, mutual information estimates give one number to characterize responses to all environmental inputs, and this analysis method characterizes how sensors respond to each environmental input. Categorical analyses, meanwhile, indicate the presence of memory without much prediction in confusion matrix elements in cultured neural networks, adding to previous understanding from mutual information estimates.

**Author summary:** All living organisms use external stimuli to navigate their environment via their sensors. Because encoding information costs energy, organisms retain only a fraction of the information received from their sensors, ideally information that maximizes their ability to remember past environmental states or predict future ones, key functions that support survival. To better understand how well sensor systems absorb stimulus information, we used stimulus-dependent Maximum Entropy (MaxEnt) models with maximum likelihood estimation and typical statistical metrics, such as confusion matrices or bias and variance. This approach provides two primary benefits over previous approaches: it is more computationally efficient, and it provides a more information-rich picture on how sensors interact with stimulus.

## Introduction

Biological organisms understand their environment and subsequently make informed decisions by using their sensors to encode information about the environment through interaction with external stimuli. This produces behavior that represents memory of past stimuli, prediction of future stimuli, or environmental conditions, posited to be the most important functions guiding how any organism navigates its environment [1, 2]. However, because encoding and transmitting information costs energy, sensory systems likely keep only a fraction of information influx from sensors, ideally information which maximizes memorization and predictive capabilities. Given this constraint, we are interested in determining how well biological organisms capture information about their environment through their sensors and to develop quantitative metrics for how stimuli are represented by biological systems.

Mutual information (MI) [3], which provides insight into how much one signal’s uncertainty is reduced by knowledge of an associated signal, has traditionally been used to gain a better understanding of the interaction between stimulus and network, such as in Ref. [4]. However, it is often difficult to calculate, with many methods running into undersampling, often with high bias and/or high variance [5], though there have been many attempts to fix these issues [6–8]. Moreover, mutual information ultimately only provides one metric about the system, thus sacrificing subtleties of how the model performs, especially when analyzing categorical variables.

An alternative method for analyzing biological systems and understanding how sensory systems interact with stimuli is via Maximum Entropy models in combination with Maximum Likelihood Estimation [9–11], quantified with typical statistics metrics. For ordinal environmental variables, e.g. where states are best considered to be real numbers or collections of real numbers, this method is more computationally efficient than MI, while the metrics it yields — bias and variance — are functions of the ordinal variable, therefore providing much more insight into the system compared to a single value from MI alone. Model performance on categorical variables, where environmental states are best considered to be discrete, meanwhile, are best quantified by confusion matrices. For *n* categorical variables, we can retrieve *n*^2^ − *n* metrics that help us quantify model performance, also exceeding the amount of information that MI provides. The extra information gives additional insight into the system, showing what kinds of input the sensor prefers and understands.

In this paper, we utilize the extra information that MaxEnt models provide and apply various versions of the model on both ordinal and categorical variables, quantifying what such models hypothesize about the integrity of environmental encoding through bias, variance, and confusion matrix elements. We start by describing experimental methods, including experimental procedure, estimator development, and statistical analysis in Materials and Methods. In the Results section, this is followed by a discussion on obtaining a certain environmental state *x* given sensor state *s* and presenting metrics for both ligand-receptor binding and memory and prediction in an organoid with commentary on observed trends. We wrap up the paper with a summary of key takeaways and directions for future work in the Conclusions section.

## Materials and methods

### Statistical mechanical models

Even though statistical mechanical models are well-founded in biophysics by physical arguments, they can be thought of as stimulus-dependent MaxEnt models. The formulation is encapsulated in biophysics textbooks [12]. Essentially, the program is that you can, like in Refs [13, 14], write down states of the system, write down multiplicities, write down energies, and from that get statistical mechanical weights and probabilities corresponding to a biological system. Weights of a state are given by the Boltzmann factor *n* exp(−*E/k*_*B*_*T*), where *n* is the multiplicity (the number of ways the state can exist) and *E* the energy of the state. The partition function *Z* is the sum of all weights, and probabilities are weights normalized by the partition function *Z*. Even though biological systems are not in equilibrium, equilibrium statistical mechanical models nonetheless serve as a good approximation [15].

### Culture preparation

We reanalyzed experimental data that were obtained from in vitro cultures of dissociated rat cortical neurons on multi electrode arrays at University of Twente, Netherlands in a study by Lamberti et al. Below we give a summary of culture preparation, recording setup, and experimental design. Readers interested in further details are encouraged to peruse Ref [3].

Neurons were obtained from newborn rats, dissociated by trypsin treatment and trituration, and then plated on multi electrode arrays (MEAs; Multi Channel Systems, Reutlingen, Germany). MEAs were stored in an incubator, under standard conditions of 36°C, high humidity, and 5% CO_2_ in air. All cultures were grown for at least 3 weeks before experiments started, to allow for culture maturation [16–18]. For experiments cultures were transferred to a recording setup in which standard conditions of 36°C, high humidity, and 5% CO_2_, were maintained. Recording began after a 15 minute accommodation period. At the end of each experiment, cultures were returned to the incubator. To make cultures responsive to light (application of optogenetic stimulation) we virally transfected them with an adeno associated virus that contained the ChannelRhodopsin-2 gene, driven by the CaMKII*α* promoter, which is found exclusively in excitatory neurons.

All surgical and experimental procedures were approved by the Dutch committee on animal use (Centrale Commissie Dierproeven; AVD110002016802) and complied with Dutch and European laws and guidelines.

### Recording setup

To record activity, MEAs were placed into a setup outside the incubator, consisting of a MC1060BC preamplifier and FA60s filter amplifier (both MultiChannelSystems GmbH, Reutlingen, Germany). Signals from the network were recorded by 59 electrodes using a custom-made Lab-View program, at a sampling frequency of 16 kHz per electrode. Acquired analogue signals were band-pass filtered (2^nd^ order Butterworth 0.1 – 6 kHz) before sampling [3, 9]. Action potentials were detected whenever recorded voltages exceeded a threshold, set at 5.5 times the estimated root mean square noise level (ranging 3 − 5 µV).

First, one hour of spontaneous activity was recorded in all cultures, followed by 20 hours of focal or global stimulation. Focal electrical stimulation was applied through one single electrode, for global (optogenetic) stimulation, power LEDs were placed above the MEA. Inter-stimulus intervals (ISIs) were drawn independently and identically from a density distribution designed to produce long-range temporal correlations (fractal renewal process) [19] and read from a pre-generated list.

### Estimate accuracy

In all cultures, action potentials were recorded from 59 electrodes, which included spontaneous activity as well as responses to optogenetic or electrical stimulation. Time stamps of detected action potentials and corresponding electrode numbers were stored and analyzed offline. All cultures were stimulated for 20 hours. Neural activity and the environmental stimulus were then binned by creating successive windows of a certain temporal width Δ*t* of activity, or bins, in which one or more spikes resulted in a 1 in that bin and no spikes resulted in a 0 in that bin. This resulted in 720, 000 total bins when assuming a time resolution of Δ*t* = 100 ms. Responses to electrical stimulation were typically recorded in the time bin of stimulation and the consecutive bin. Due to the relatively slow opening of ion channels upon light activation, responses to optogenetic stimulation reached their maximum within the two bins following the stimulus bin.

Activity and stimulation in each bin is represented by a 60 element vector (59 electrodes and 1 stimulus), thus creating a 60 by 720, 000 binary matrix. This matrix contains mostly zeros, as only about 15, 500 of the 720, 000 bins (2.15%) of the bins contain stimulation, and most of the electrodes are inactive in the majority of bins, particularly when not stimulated.

Due to computational complexities discussed thoroughly in the “Applying the model” subsection of the Results section, a subset (ranging between three and six electrodes) of the prepared binary matrix was analyzed using the Maximum Entropy spin-glass Ising model to estimate *J*_*xx*_, *J*_*xs*_, *J*_*sx*_, *J*_*ss*_ in Eq 6 and subsequently to calculate a likelihood ratio *L* (Eq 24) and estimator 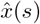. Finally, 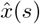 was compared to the stimulus row list of the real data *x*. Each match between 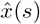 and *x* was summed and divided by the amount of total bins to compute an overall estimate accuracy of the experiment, and a confusion matrix was generated tabulating the errors made in guessing *x*.

### Statistical Analysis

The Shapiro-Wilk test was used to confirm normality within treatments. On the condition of normality, *t*-tests were used to determine the significance of differences between the control and stimulated experiments. Data were alternatively analyzed with nonparametric tests if the Shapiro-Wilk test determined a non-normal distribution. Statistical significance is determined using an *α*-level of *α* = 0.05.

## Results

### Stimulus-dependent MaxEnt models as sensor models

In order to proceed, we need *p*(*x*|**s**), where *x* is the environmental variable and **s** is the sensor state. This is achieved either through a stimulus-dependent MaxEnt model or through an equilibrium statistical mechanical model that incorporates an environmental variable.

#### Stimulus-dependent MaxEnt models

To make a stimulus-dependent MaxEnt model, we depart somewhat from Refs [10, 11] and instead make a MaxEnt model of the joint probability distribution of environment *x* and sensor **s**. These models take the form of an exponential linear model,

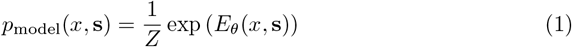

regardless of constraints used with parameters *θ* used to fit the data distribution so that *D*_KL_[*p*_data_||*p*_model_] is minimized, where

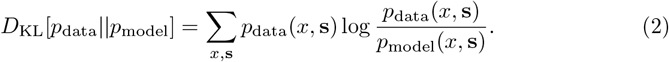

This is an alternative formulation of MaxEnt models that differs from the usual treatments in which we aim to make a model that is as unbiased as possible using given information from data, such as in Ref [10]. From the joint MaxEnt model one can derive

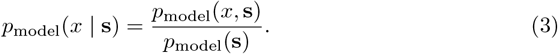

It turns out that for the analysis described later, we will not need *p*_model_(**s**). As such, the joint MaxEnt model is as good as the stimulus-dependent MaxEnt model [11] for sensor analysis. If instead *p*_model_(**s**|*x*) is obtained, *p*(*x*|**s**) can be approximated using Bayes rule, as we describe later in Eq 8.

Here we give a brief summary of Ref [10] to illustrate the typical MaxEnt model. Neuronal activity between time *t* and *t* + Δ*t* is encoded by a binary vector ***σ***, where Δ*t* = 100 ms. If neuron *i* fires in that time frame, *σ*_*i*_ is set to 1, otherwise, *σ*_*i*_ = 0. After construction of all binary vectors that occur during a long term recording, their probability distributions are modeled as

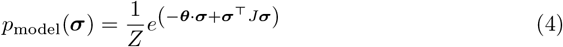

where *Z*, the partition function, is a normalization factor:

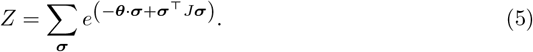

In Eq 5, ***θ*** is a vector that represents the propensity of neurons to fire. *J* is symmetric by definition, and represents first order interactions between neurons.

To adapt such a model for our use case, we fit a joint MaxEnt model to the stimulus *x* and network activity **s**, *p*_model_(*x*, **s**). This is done by concatenating the network activity vector **s** and stimulus *x* into a binary vector ***σ*** whose first elements describe neural activity and whose last element indicates whether or not there was a stimulus. We can “time-shift” *x* from **s** to simulate memory or prediction, as demonstrated in Ref [3].

We thus consider the vector 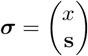 and maximize the entropy of *p*(***σ***) while constraining the mean of ***σ***, ⟨***σ***⟩, and the second moment, ⟨***σ*** · ***σ***⟩. This leads to the joint MaxEnt model

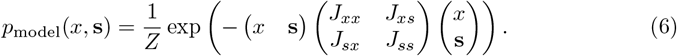

Eq 6 incorporates both constraints: because ***σ*** is a binary vector, each element satisfies 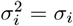, and therefore the diagonal of *J* accounts for the mean constraint by doubling as a magnetic field analogue from the Ising model.

#### Statistical mechanical models

Statistical mechanical models are, in some way, examples of stimulus-dependent MaxEnt models. An example is shown in Fig 1 for the lac repressor genetic regulatory circuit described in Ref. [13]. The result of this statistical mechanical calculation is the conditional probability of RNAP binding to the promoter region given that there is a certain number of lac repressor molecules. We may use this variable as the environmental variable due to the intimate relationship between it and lactose molecules: as lactose molecules — the true environmental variable — appear, they bind to lac repressor molecules in a chemical reaction that leads to a certain concentration of free lac repressor molecules being a direct readout of the concentration of lactose molecules. The output of the statistical mechanical model [13] is the following model for *p*_model_(*s* | *x*), where *s* is whether or not RNAP is bound and *x* is the number of lacR molecules:

**Fig 1.**
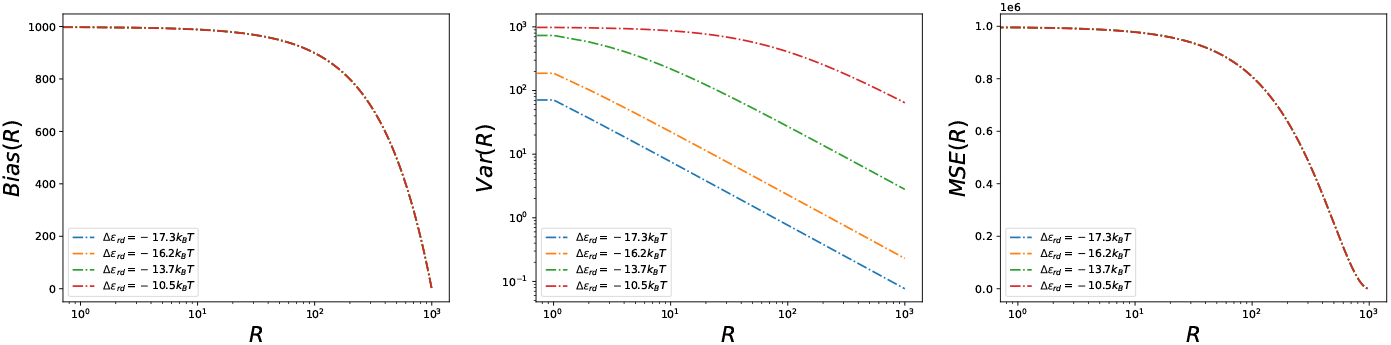
Bias, variance, and MSE of the lacR estimator based on whether or not RNAP is bound. At left, bias as a function of the number of lacR molecules *R*. In the middle, variance as a function of the number of lacR molecules *R*. At right, mean-squared error (MSE) as a function of the number of lacR molecules *R*. The energy of lacR binding is varied and *N*_*NS*_ = 5 × 10^6^ chosen as in Ref [13].

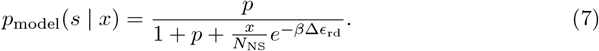

Here, *p* ≈ 10^−3^ and *N*_NS_ ≈ 5 × 10^6^, whereas Δ*ϵ*_rd_ adopts a range of values that can be tuned in experiments [13].

With *p*(*s* | *x*), we can find *p*(*x* | *s*) via Bayes’ theorem:

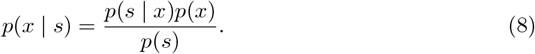

We will often assume a uniform input concentration *p*(*x*), as it is the most unbiased model of the environment with no constraints. The sensor marginal distribution *p*(*s*) will not be explicitly needed, as we will see in the next subsection.

Another example of a statistical mechanical model that describes everything from bacterial chemotactic receptors to nACh receptors is the Monod-Wyman-Changeux (MWC) model [14], according to which the probability of a receptor with *n* sites being active is

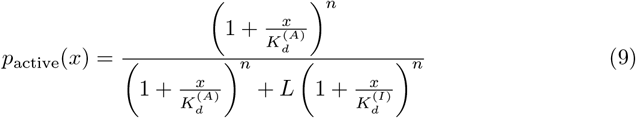

for dissociation constants 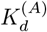 and 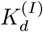 and a parameter *L* that governs the energetic difference between the active and inactive state. This is alternatively *p*(*s* | *x*), where *x* is the ligand concentration and *s* is whether or not the MWC molecule is active.

### Analyzable estimators of the environment from sensor models

Once we have *p*_model_(*x* | **s**), we utilize Maximum Likelihood Estimation to form an estimator of the environment from the sensor state:

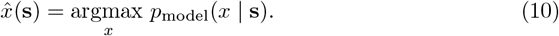

If we have a joint MaxEnt model, we can find this as

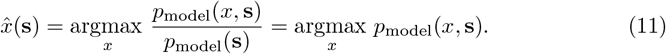

If instead we have *p*_model_(**s** | *x*) as with the statistical mechanical models, we can find the estimator as

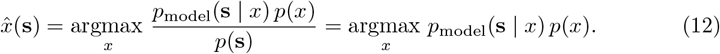

We can approximate *p*(*x*) from data or from another model. For our particular neural system, we use *p*(*x*) = *p*_data_(*x*). We could instead use an unbiased model of the environment, making *p*(*x*) uniform, leading to

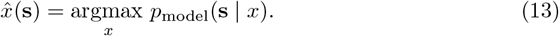

In making these estimates, we imagine a downstream region of the brain or organism finding the sensor state and trying to invert it to find an estimate of the environment. This estimate of the environmental state can then be used to inform action policy.

### Bias, variance, and mean-squared error with ligand-receptor binding

If the environmental variable is ordinal, then we can form bias, variance, and mean-squared error estimates of the environmental variable as follows:

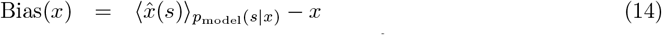

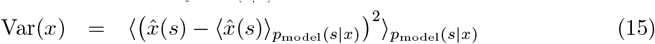

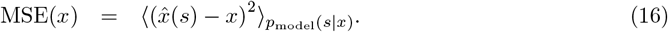

We can use MSE(*x*) = Bias(*x*)^2^ + Var(*x*) to calculate MSE easily. Bias, variance, and MSE of an estimator are all indicators of the estimator’s quality. Bias indicates the accuracy, while variance indicates its precision and MSE indicates its overall quality.

To illustrate, bias, variance, and MSE are calculated for the simple genetic regulatory circuit in Ref [13] in Fig 1 below. We find that when RNAP is bound, we should estimate that there are no lacR molecules in the environment, and that when RNAP is not bound, we should estimate that there are as many lacR molecules in the environment as possible, regardless of binding energy Δ*ϵ*_rd_. Typically, RNAP is not bound. As such, this estimator leads to bias estimates that are large when the number of lacR molecules is small and MSE estimates that are large when the number of lacR molecules is small. The MSE is mostly dominated by the bias. The variance (estimator noisiness) depends more intricately on the details of the binding energy of lacR to the DNA site. When the binding energy is less favorable, we see a noisier estimator of lacR molecules, as the probability of RNAP binding is higher overall.

Bias, variance, and MSE are also calculated for the nACh molecule using the Monod-Wyman-Changeux model from parameters in Ref [20] and shown in Fig 2. The estimator gives a maximal ACh concentration estimate when the nACh receptor is active (open) and an ACh concentration estimate of 0 *M* when the nACh receptor is inactive (closed). From this, bias is maximal at middling concentrations. Variance is maximized when the probability of the receptor being active is 1*/*2. Together, these determine a mean-squared error estimate that is large at middling concentrations, due to large bias and variance. Unlike the channel capacity calculation in Refs [14, 21], we do not need to assume that the organism controls the environment (*p*([ACh])), and we also gain insight into which environmental variables are well-represented– the analogue of understanding the optimal *p*([ACh]) that maximizes mutual information in Refs [14, 21], which shows peaks at low and high concentrations.

**Fig 2.**
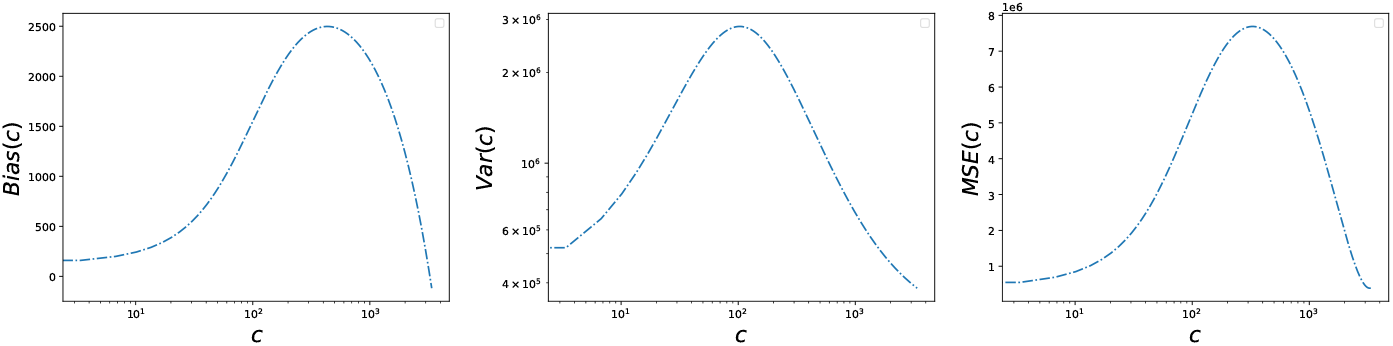
Bias, variance, and MSE of the ACh concentration estimator based on whether or not nACh is in the active (open) state. At left, bias as a function of the ACh concentration *c*. In the middle, variance as a function of the ACh concentration *c*. At right, mean-squared error (MSE) as a function of the ACh concentration *c*. Parameters are taken from Ref [20].

Bias, variance, and MSE are also calculated for the bacterial chemotactic receptors using the Monod-Wyman-Changeux model from parameters in Ref [22] and shown in Fig 3. The estimator gives a minimal chemoattractant concentration estimate when the bacterial chemotactic receptors are inactive and a high but not maximal chemoattractant concentration estimate when the receptors are inactive. Again, from this, bias is maximal at middling concentrations and at higher concentrations than the maximal concentration estimate. Again, variance is maximized when the probability of the receptor being active is 1*/*2. Together, these determine a mean-squared error estimate that is small at smaller concentrations and larger at larger concentrations, with a minimum at the maximal concentration estimate. Unlike a channel capacity calculation, we do not need to assume that the bacterium controls the environment (*p*(*x*)), and we also gain insight into which environmental variables are well-represented– the analogue of understanding the optimal *p*(*x*) that maximizes mutual information, which shows valleys when *variance* of the estimator is low. This calculation reveals not only the noisiness of the estimator but also the accuracy of the estimator from the bias, and therefore provides more detail than the corresponding channel capacity calculation.

**Fig 3.**
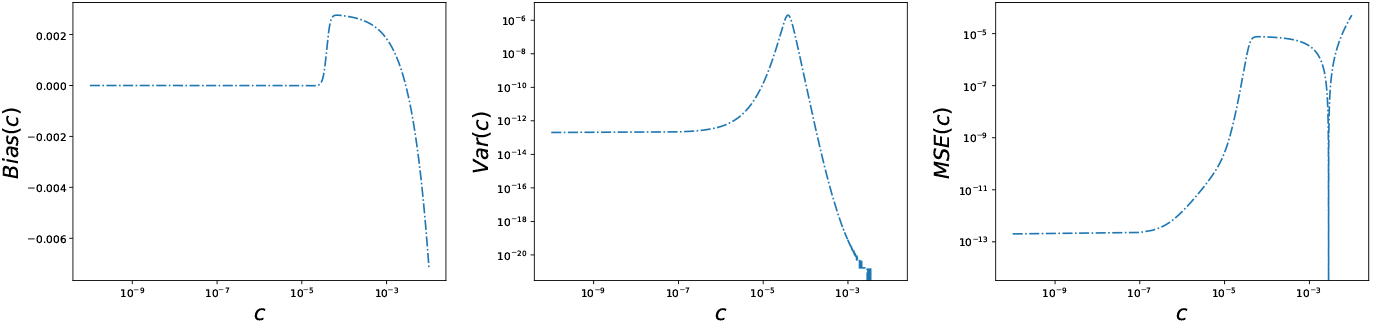
Bias, variance, and MSE of the chemoattractant concentration based on whether or not the receptors are in the active state. At left, bias as a function of the chemoattractant concentration *c*. In the middle, variance as a function of the chemoattractant concentration *c*. At right, mean-squared error (MSE) as a function of the chemoattractant concentration *c*. Parameters are taken from Ref [22].

To showcase new insights that can be obtained using this analysis method, we turn to asking for the benefit or downside of cooperative sensors. Previously, cooperativity in sensors was shown to aid information processing [14, 21, 23]. Yet, a simple mutual information analysis says independent is better: *I*[*s*_1_, *s*_2_; *x*] = *I*[*s*_1_; *x*] + *I*[*s*_2_; *x* | *s*_1_] is maximized if *s*_1_, *s*_2_ are conditionally independent given *x*. Does cooperativity in a two-state Adair model, described below, do anything to information processing as measured by bias, variance, and MSE? We have:

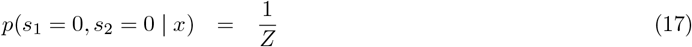

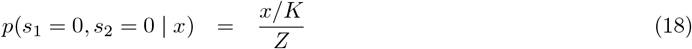

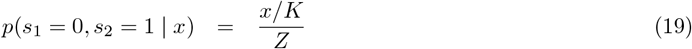

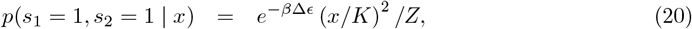

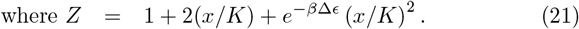

If we look at MSE, there is a range of ligand concentrations for which cooperativity improves performance (lower MSE) and a range of ligand concentrations for which it does not. The answer to the question of whether or not cooperativity benefits sensor performance depends on variables such as what kinds of ligand concentrations the sensor is likely to see. See Fig 4.

**Fig 4.**
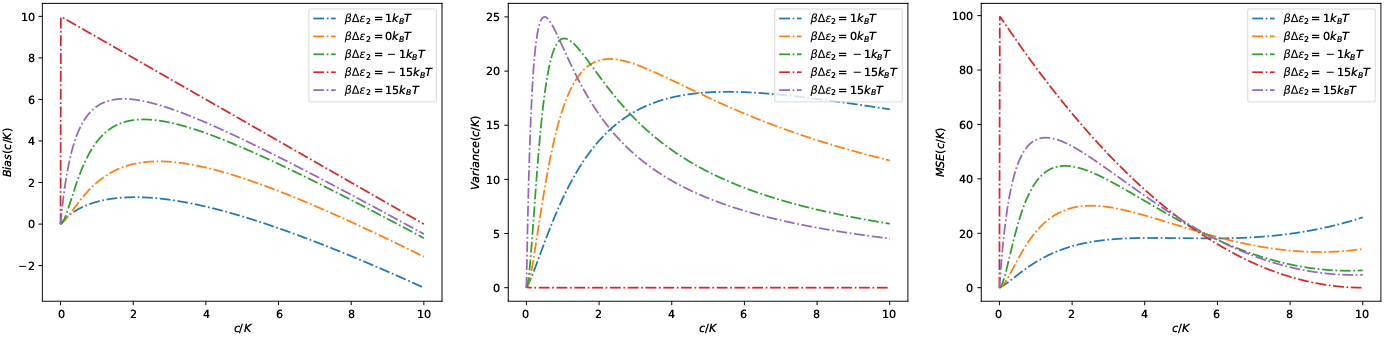
The bias, variance, and MSE of a sensor with varying degrees of cooperativity. At left, bias as a function of the chemoattractant concentration *c*. In the middle, variance as a function of the chemoattractant concentration *c*. At right, mean-squared error (MSE) as a function of the chemoattractant concentration *c*. A range of cooperativities based on varying *β*Δ*ϵ* are shown.

### Confusion matrices with memory and prediction in cultured neurons

Because behavior in all cultures was monitored by 59 electrodes [3], whose signals were used to construct a prediction about the activity of the optogenetic or electrical stimulus, our estimator ultimately makes a categorical prediction. Therefore, it is most productive to use confusion matrices to quantify model performance. Since the estimator guesses the activity or inactivity of the stimulus, it is binary, and thus we may avoid calculating the partition function and instead develop a likelihood ratio

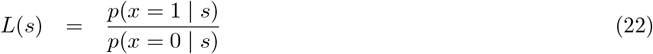

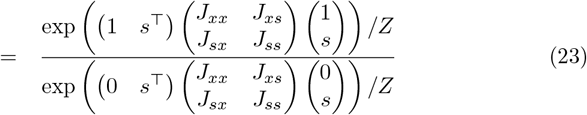

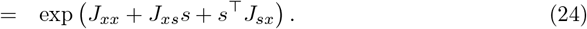

If *L*(*s*) > 1, then 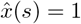; otherwise 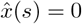.

#### Creating controls

When assuming a time bin of Δ*t* = 100 ms, approximately 97.85% of all entries in the stimulus electrode were inactive, thus positively biasing the accuracy of stimulus estimates 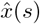. To assess this bias and its effects on confusion matrix outputs, control data were generated and analyzed using the MaxEnt model. First, assuming no correlation between any of the neurons and a random firing rate, completely random data are generated, and the estimator should not be biased into guessing either activity or inactivity over the other. Thus, the estimator should predict correctly half the time, resulting in a confusion matrix as in Table 1.

**Table 1.**
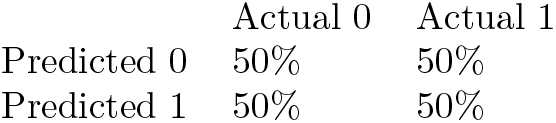
The confusion matrix generated for completely random data. Given this assumption, the model is expected to correctly predict both electrode activity (true positive rate, bottom right entry) and inactivity (true negative rate, top left entry) half of the time. Incorrect predictions are documented in off-diagonal entries, and each column must sum to 100% to account for all data.

|  | Actual 0 | Actual 1 |
| --- | --- | --- |
| Predicted 0 | 50% | 50% |
| Predicted 1 | 50% | 50% |

Next, we relax the assumption that none of the neurons are correlated but maintain the assumption that the stimulus and network are uncorrelated. Under this constraint, the off-diagonal terms in the *J* matrix — related to the correlation strengths between neurons [9, 10] — do not contribute to the likelihood ratio, which thus is dominated by the activity propensity of the stimulus. To generate simulated data under these conditions, we constructed a purely diagonal *J* matrix with negative coefficients in the diagonal to reflect the fact that electrodes are more likely to be inactive than active, then analyzed it using our model. The model defaults to estimating exclusively inactivity for the duration of our experiment, thus achieving 97.85% baseline accuracy at a time bin of Δ*t* = 100 ms.

#### Applying the model

To develop an estimator, we used the Maximum Entropy spin-glass Ising model. Because this model requires calculation of the partition function, we chose a subset of the 60 neurons to construct the network vector *s*. Calculating the full partition function for *n* neurons requires summing over all 2^*n*^ configurations of the system and, for our case of 60 neurons, is not computationally feasible. Therefore, it is worth investigating how much information can be captured with a subset of the entire population. To choose a subset, a so-called correlation “vector” was constructed between the stimulus and each neuron in the network using Pearson correlation coefficients, from which the *n* network neurons with the highest correlation to the stimulus were chosen for a total system size of dimension *n* + 1. In this case, the *J* matrix is a square matrix with (*n* + 1) rows and columns. Subsystem sizes were chosen from three to six neurons by choosing the neurons with highest Pearson correlation coefficients. The MaxEnt model was then fitted to the data, and the confusion matrix metrics characterizing the estimator’s performance are shown in Fig 5.

**Fig 5.**
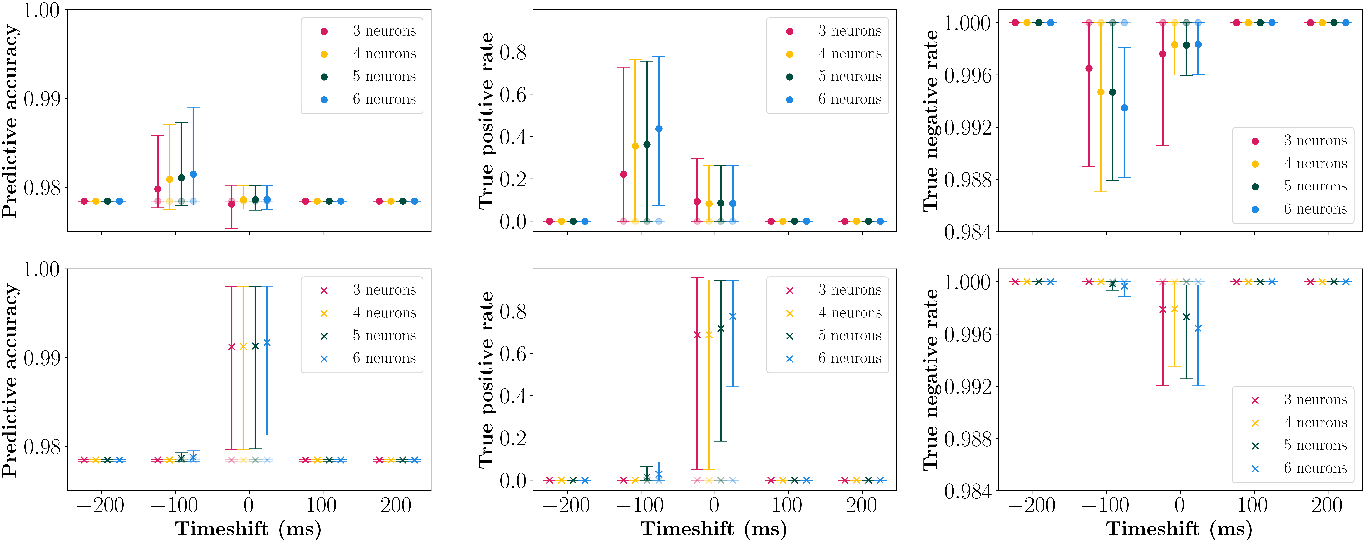
Predictive accuracy, true positive rate, and true negative rate for baseline and model estimators. Colors correspond to varying neuron subsystem sizes used for determining the *J* matrix. Circular markers denote optogenetic (*n* = 10) stimulation (top row), crossed markers denote electrical (*n* = 8) stimulation (bottom row). 90% confidence intervals are constructed by marking the 5th and 95th percentile of each metric over all experiments. Plots with varying system sizes at the same time-shift are staggered to enhance visibility of markers and error bars.

This model can also be applied after time-shifting the stimulus *x* relative to the network activity *s*, allowing us to assess prediction and memory confusion matrices. Incorporating a positive-signed time-shift is physically interpreted as assessing the system’s prediction capability, while a negative-signed time-shift assesses the system’s memory capability [3]. Time-shifting was performed out to ±200 ms; larger time-shifts relegated model performance to baseline.

Statistics of model metrics over experiments were explored to compare optogenetic and electrical trials. At a timeshift of 0 ms and a subsystem size of six neurons, both local and global model predictions were found to be distributed normally. With *α* = .05, performing a Shapiro-Wilk normality test on the aforementioned sets yielded *p* = 0.0897 and *p* = 0.194 for local and global trials, respectively. Moreover, we found a significant difference in predictive accuracies between globally and focally stimulated cultures, where focally stimulated cultures showed a 1.31% improvement in performance with an associated *p*-value of *p* = 1.47 · 10^−5^. Meanwhile, there was no statistically significant difference between globally stimulated cultures and control trials (*p* = 0.524).

Examining electrical stimulation, model performance most significantly improves over the baseline when there is no time-shift with a mean model predictive accuracy at 99.14% and true positive rate of 71.35%. Simultaneously, zero time-shift also presents the most significant drop in true negative rates, with correct predictions occurring 99.75% of the time. Fig 5 shows an asymmetry in time-shifting 100 ms, with an incremental improvement in model predictive accuracy compared to baseline for −100 ms and no improvement in the positive direction.

Applying the model to optogenetic stimulation experiments yields most significant improvements at −100 ms time-shift, with a mean model predictive accuracy of 98.08%, true positive rate of 34.41%, and true negative rate of 99.48%. At 0 ms time-shift, marginal improvements in true positive rate are observed (about 8.61%), but the decrease in true negative rate negates model predictive accuracy to be comparable to baseline.

Fig 1d in Ref [3] provides more insight to the observed trends. Mean number of spikes is highest immediately following stimulus activity in electrical stimulation, with a vanishing minority of spikes occurring between 100 and 200 ms post-activity. This translates to the system’s memory of the stimulus being strongest right after stimulus activity, which shows up as a significant improvement in model predictive accuracy at 0 ms time-shift. The distribution in electrode spikes for optogenetic stimulation, however, is notably wider because the light pulse itself lasts about 100 ms. Therefore, the mean number of spikes is densest at around 100 ms post-activity. Network electrodes for optogenetic stimulation also do not start spiking until several milliseconds post-activity as opposed to almost immediately for its electrical counterpart, which further accounts for the drastic observed difference in predictive accuracy at 0 ms time-shift.

Varying electrode subset sizes also reveals interesting behavior. Remarkably, even selecting the two network electrodes with highest correlation with the stimulus captures a majority of the improvements in predictive accuracy, evidenced by marginal improvements with increasing system size. While, in general, the predictive accuracy shows marginal improvement with respect to system size, certain trials exhibit counterintuitive behavior, where increasing system size does not imply a monotonically non-decreasing predictive accuracy.

Moreover, regardless of stimulation type, none of the systems explicitly displayed any predictive behavior, since at any time-shift +100 ms and beyond, all model confusion matrix metrics were identical to baseline. Ref [3], however, concludes that the same system possesses predictive capability dependent on the system’s short and long-term memory. This discrepancy most likely comes down to the choice of selecting system subsets. Subsets in the MaxEnt model were chosen based solely on correlation coefficients examined holistically over the duration of the experiment, not in an attempt to maximize mutual information between stimulus and network. While the chosen electrodes are most strongly correlated to the stimulus and provide a preliminary guide to choosing an optimal subset for developing the estimator, the correlation coefficients remove time dependence in its analysis, and thus likely fails to capture network dynamics that tease out more information about the stimulus, including patterns demonstrating stimulus prediction. Alternatively, we could have added neurons to the subsystem that maximally improved predictive accuracy as in Ref. [3], but doing so requires an exhaustive search through all neurons, which is more computationally expensive than our current approach.

To understand the effects of time resolution on model metrics, we additionally ran the MaxEnt model on our experiments while varying time resolution, using time bins of Δ*t* ∈ {25, 50, 100, 200} ms. We used a subsystem size of six electrodes at 0 ms time-shift for electrical stimulation and −100 ms time-shift for optogenetic stimulation to best investigate how each confusion matrix metric varied with respect to experimental time resolution.

While the leftmost subfigure in Fig 6 does not show a statistically significant improvement in predictive accuracy improvement against baseline with respect to time resolution coarse-graining, we can gain insight into observed mean true positive and negative rate trends. Mean true negative rate monotonically decreases for both electrical and optogenetic stimulation, while mean true positive rates tend to increase, with the exception of optogenetic stimulation at 200 ms time bins.

**Fig 6.**
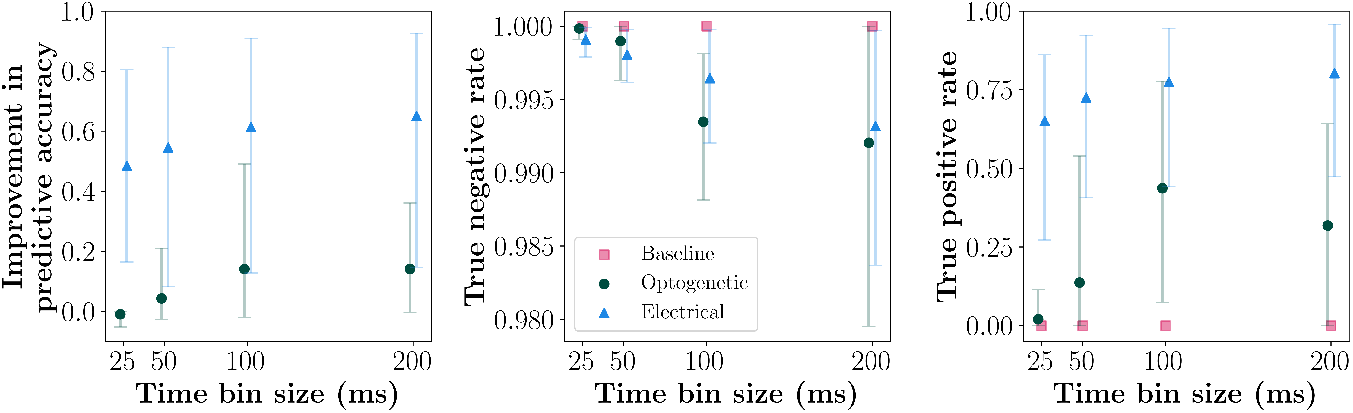
Improvement in predictive accuracy relative to baseline (where 1.0 represents perfect predictive accuracy), true positive rate, and true negative rate for baseline and model estimators. Colors and marker types correspond to stimulation type (red is baseline) used for determining the *J* matrix. Circular markers denote optogenetic (*n* = 10) stimulation and constitute the first row, while crossed markers denote electrical (*n* = 8) stimulation and constitute the second row. 90% confidence intervals are constructed by marking the 5th and 95th percentile of each metric over all experiments. Plots with varying stimulation type at the same time resolution are staggered to enhance visibility of markers and error bars.

At finer time resolutions, network patterns that otherwise indicate association with the stimulus are divided into multiple time bins, resulting in fewer instances where the model can pick up on network behavior and correctly guess stimulus activity. This is much less of a problem for electrical stimulation, since most electrode spikes for the former stimulation type occur within a few milliseconds of stimulus activity (Fig 1d of Ref [3]), and therefore spike patterns of electrodes most correlated to the stimulus are not significantly disturbed even at a time resolution of Δ*t* = 25 ms. Meanwhile, in the optogenetic case, where most spikes are recorded about 100 ms after stimulus activity, improvements in the true positive rate are most pronounced as time resolution are coarse-grained up to Δ*t* = 100 ms, and network spike patterns emerge as previously separate time bins are fused. Simultaneously, true negative rate is hypothesized to decrease as time bin size increases because network spike patterns that correspond to stimulus inactivity are combined with those that correspond to stimulus activity, consequently resulting in more false positives. Additionally, if time resolution is too coarse, the system loses network spike patterns that differentiate whether the stimulus is active and choosing electrodes most correlated to the stimulus becomes increasingly challenging, ultimately resulting in poorer model performance.

Finally, experiments were decomposed into hour-by-hour segments to investigate whether there were incremental differences throughout the duration of stimulation. Inspecting Fig 7, regardless of stimulation type, there was slight fluctuation from hour to hour but no significant increase or decrease of prediction ability over the course of a 20 hour experiment. The observed high correlation between uncorrelated baseline and global stimulation in Fig 7, while remarkable, is explained by the lack of statistical significance between globally stimulated cultures and control trials. Such results, combined with no observable predictive capability, suggest that the network population may not learn the stimulus but nonetheless responds to stimulus activity via reverberation of the stimulus around the network.

**Fig 7.**
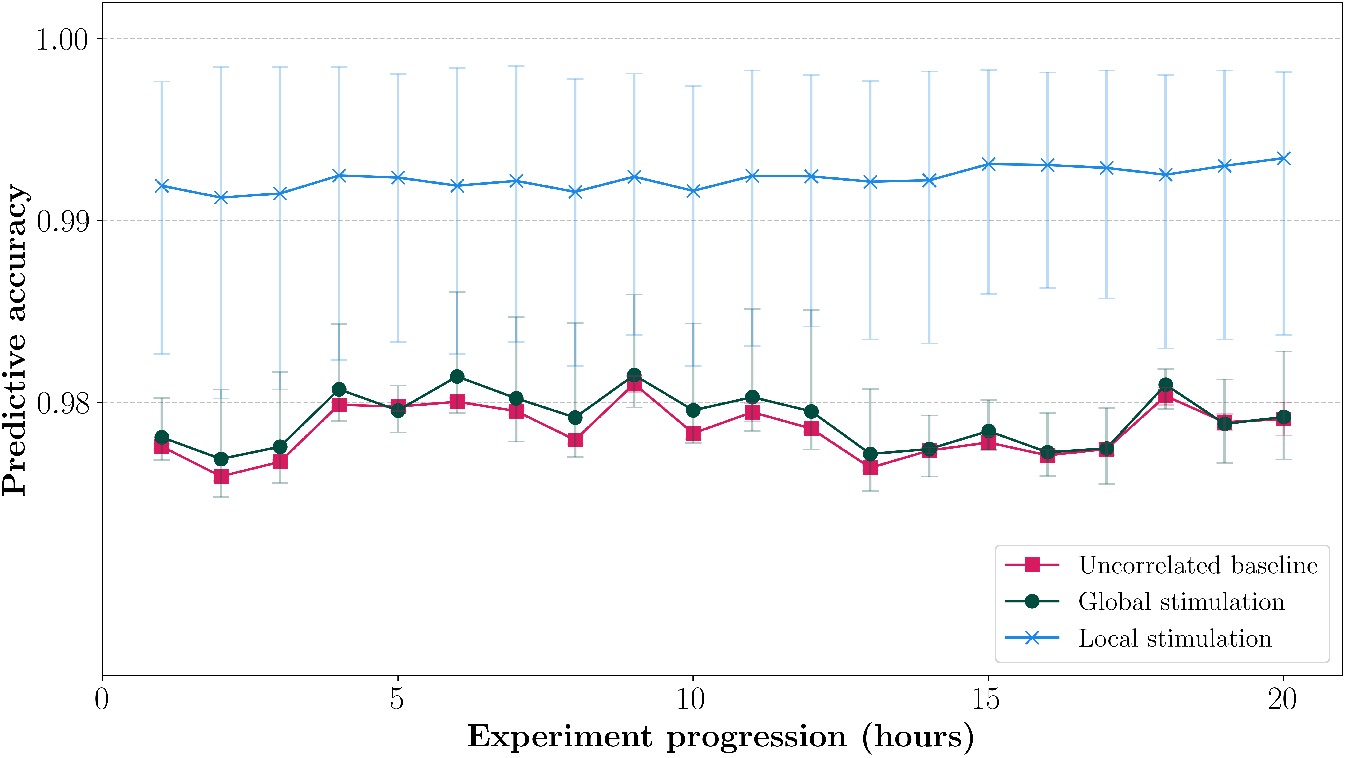
Hour-by-hour mean MaxEnt model predictive accuracy for optogenetic (*n* = 10) and electric (*n* = 8) stimulation experiments for the experimental duration (20 hours) relative to an uncorrelated baseline estimate. Accuracy was determined with a subsystem of six electrodes at 0 ms time-shift. Error bars represent a 90% confidence interval.

## Conclusion

We applied and investigated a combination of stimulus-dependent Maximum Entropy models [11] or joint Maximum Entropy models with maximum likelihood estimation and the usual statistical metrics on both ordinal and categorical variables to understand how well sensors represent environmental information. These methods use the traditional Maximum Entropy models [10, 11] to understand how a homunculus might see the environment to provide an information-rich alternative to mutual information assessments. This method reveals not only that sensors represent some amount of information about the environment, but how much this amount of information varies and what type of information is provided based on the environmental input.

When the environmental input was ligand concentration, the analysis revealed how much bias and variance was present in the sensory estimator’s implicit understanding of the environment. We analyzed a number of MWC receptors as in Ref [21] and found new insights into exactly how well these receptors represented their ligand concentrations. In particular, we found that while a mutual information analysis shows that independent receptors without cooperativity transmit more information about the environmental input, the analyses here showed that some ligand concentrations were better estimated with cooperative receptors based on bias and variance metrics.

We looked at cultured neurons stimulated by a point process stimulus and analyzed its memory and predictive power with this new analysis as opposed to the mutual information analysis in Ref [3]. We treated the joint stimulus and sensor with a spin-glass Ising model to quantify memory and prediction via confusion matrices, with true positive, true negative, and predictive accuracies visualized as a function of time-shift, subsystem size, and stimulation type. These metrics yielded results consistent with post-stimulus electrode spiking distributions observed in Ref [3] but additionally quantified how strongly the network encodes memory of the stimulus through illustrating exact numerical changes in true positive and negative rates with time-shift. Moreover, increasing subsystem size reveals improvements in average predictive accuracy of the network, which has not been thoroughly investigated yet. Changing time resolution showed a time bin size coarse enough to group stimulus activity indicators yet fine enough not to wash out system details yielded the best estimators, while partitioning experiments into hourly intervals showed that networks represent the stimulus in a way that is stable over the entire experiment.

Mutual information is notoriously difficult to estimate [5, 24]. Compared to the mutual information approach [3], we faced no computational difficulties when calculating ordinal quantities with ligand-receptor binding, but did face computational difficulties in developing estimators for categorical variables. Computational difficulties primarily stemmed from calculating the partition function, whose computational complexity scales exponentially with system size. Estimating confusion matrix metrics using Minimum Probability Flow [25] or the closely-related Minimum Energy Flow [26] — a more compute-friendly parameter estimation method — was also tried, but ultimately was not fruitful because it failed to correctly predict the functional connectivity between neurons and stimulus, despite Ref [9]. Contrastive divergence is a possible workaround to such problems [27].

Despite computational difficulties in estimating stimulus-dependent [11] or joint Maximum Entropy models, our methodology nonetheless provide invaluable insights into memory and prediction in these biological systems. Mutual information analyses produce only one number to characterize the sensor’s entire performance, while our analysis reveals detailed information about how well the sensor deals with each type of environmental stimulus. Perhaps future efforts could focus on remedying this problem with mutual information analyses via a calculation of *I*[*S*; *X* = *x*], but even this fails to understand the benefits of cooperativity in receptors, for instance. As it stands, our new method is computationally tractable and produces additional insight into biological systems that would be impossible to glean otherwise.

## Supporting information

**S1 Code Repository. [Repository for confusion matrices with memory and prediction](https://github.com/chrisw47/maxent-mempred-confusionmatrix)**. This contains the code used for analysis and figure generation in the confusion matrices section of the paper. Fig 5 is generated by cm metrics plot.py, Fig 6 by binsizes.py, and Fig 7 by hourly.py. More information can be found in the README.md file located within the repository.

## Acknowledgments

This work was performed in part at the Aspen Center for Physics and was supported by a grant from the Alfred P. Sloan Foundation (G-2024-22395). Many thanks to the Pioneer Academy for support and hospitality. This study was supported by the US Air Force Office for Scientific Research, Grant Number FA9550-19-1-0411.

## Notes

### Competing Interest Statement

The authors have declared no competing interest.

